# Ionic strength modulates structural disorder and protein oligomerization in the marginally disordered Phd transcription factor

**DOI:** 10.64898/2026.04.15.718675

**Authors:** Uroš Zavrtanik, Gopinath Muruganandam, Maruša Prolič-Kalinšek, Dietmar Hammerschmid, Frank Sobott, Alexander N Volkov, Remy Loris, San Hadži

**Author notes:** These authors contributed equally to this work (Uroš Zavrtanik, Gopinath Muruganandam).

## Abstract

Some proteins combine sequence features that are typical both for folded proteins and intrinsically disordered proteins (IDPs). The borderline properties of these so-called “marginal” IDPs render their conformational ensembles highly sensitive to the environmental changes, which may be important for their function. Here, we investigate the prokaryotic transcription factor Phd, which regulates the *phd-doc* toxin-antitoxin module through an allosteric mechanism involving disorder-order transition. Using an ensemble of biophysical techniques, we show that the protein is completely disordered at low ionic strength, whereas increasing salt concentration promotes its collapse into a partially ordered monomeric state, followed by the formation of a structured dimer. Using a thermodynamic model, we decipher the linkage between ionic strength, protein stability, oligomer state and degree of disorder. Via small-angle X-ray scattering we derive the structural ensemble of dimeric Phd, revealing a gradation of disorder as a function of salt. The sequence and biophysical properties of Phd position it at the boundary between macroscopically distinct conformational ensembles, representing a large pool of states capable of engaging in functional disorder-to-order interactions, enabling Phd to act as conformational rheostat. Together with previous crystallographic data, this charts the full spectrum of disorder-to-order states in the bacterial transcription factor and underscores the structural plasticity of IDPs with the marginal sequence properties.

**SIGNIFICANCE:** While globular proteins adopt stable three-dimensional structures, intrinsically disordered proteins (IDPs) remain flexible and dynamically sample partially ordered states. This behavior is largely determined by amino acid composition; however, some proteins lie at the borderline between order and disorder. Here, we focus on such a protein, the Phd transcription, and show how its conformation changes from completely disordered to fully ordered. These transitions are modulated by ionic strength and binding to macromolecules, including homodimerization. Using a thermodynamic model, we map the Phd conformational space, revealing a broad ensemble of states with varying degrees of disorder. The borderline amino acids properties enable Phd to function as a conformational rheostat, coupling functional interactions to the series of graded order–disorder transitions.

## INTRODUCTION

Protein structures range from a very narrow structural ensemble in their folded state to highly disordered intrinsically unfolded ensembles that do not adopt a folded structure, not even when they form specific complexes with other macromolecules (for a comprehensive overview, see van der Lee *et al.,* 2014). Such intrinsically disordered proteins (IDPs) have been observed to carry out many specific functions including acting as chaperones, molecular clocks (Kovacs & Tompa, 2012; Liebovitch *et al*., 1992; Hoshi *et al*., 1990), entropic springs (e.g. in Titin - Linke *et al*., 1998), parts of signaling machinery (Wright & Dyson, 2015) and most prominently formation of biomolecular condensates (Shin & Brangwynne, 2017). IDPs are highly enriched in transcription factors: 83 to 94% of eukaryotic transcription factors possess extended intrinsically disordered regions (IDR) and intrinsic disorder is also prominent in many other proteins that are part of the transcription machinery (for a review, see Fuxreiter *et al*., 2008).

Understanding sequence features that determine the extent of order/disorder in proteins has been a key challenge in the field of IDPs. One early observation noted that, relative to globular proteins, the disordered proteins are significantly depleted in hydrophobic residues and enriched in charged ones (Uversky *et al*., 2000). This observation led to the empirical boundary between disordered and folded proteins separating these two groups on the charge-hydropathy plot (**Figure 1**). Differences in the composition of specific amino acids between disordered and globular proteins, such as enrichment of Pro and charged residues IDPs, were the basis for development of accurate algorithms that predict protein disorder from the amino acid sequence (Mészáros *et al*., 2018; Romero *et al*., 2001). In the case of charged IDPs and polyampholytes (IDPs with similar numbers of positively and negatively charged residues) metrics such as net charge per residue, fraction of charged residues and pattering of charged residues are instrumental for understanding the degree of compaction/extension of IDP ensembles (Das & Pappu, 2013; Mao *et al*., 2010; Sawle & Ghosh, 2015). An interesting subgroup are IDPs with borderline sequence properties, which are located at the folded/disordered or collapsed/expanded boundaries of the Uversky and Das-Pappu plots, respectively. It has been hypothesized that the structural ensembles of these “marginal” IDPs could be much more responsive with respect to perturbations, such as changes in solution conditions, posttranslational modifications or interactions with other molecules (Fuxreiter, 2024; Firouzbakht *et al*., 2024)

**Figure 1.**
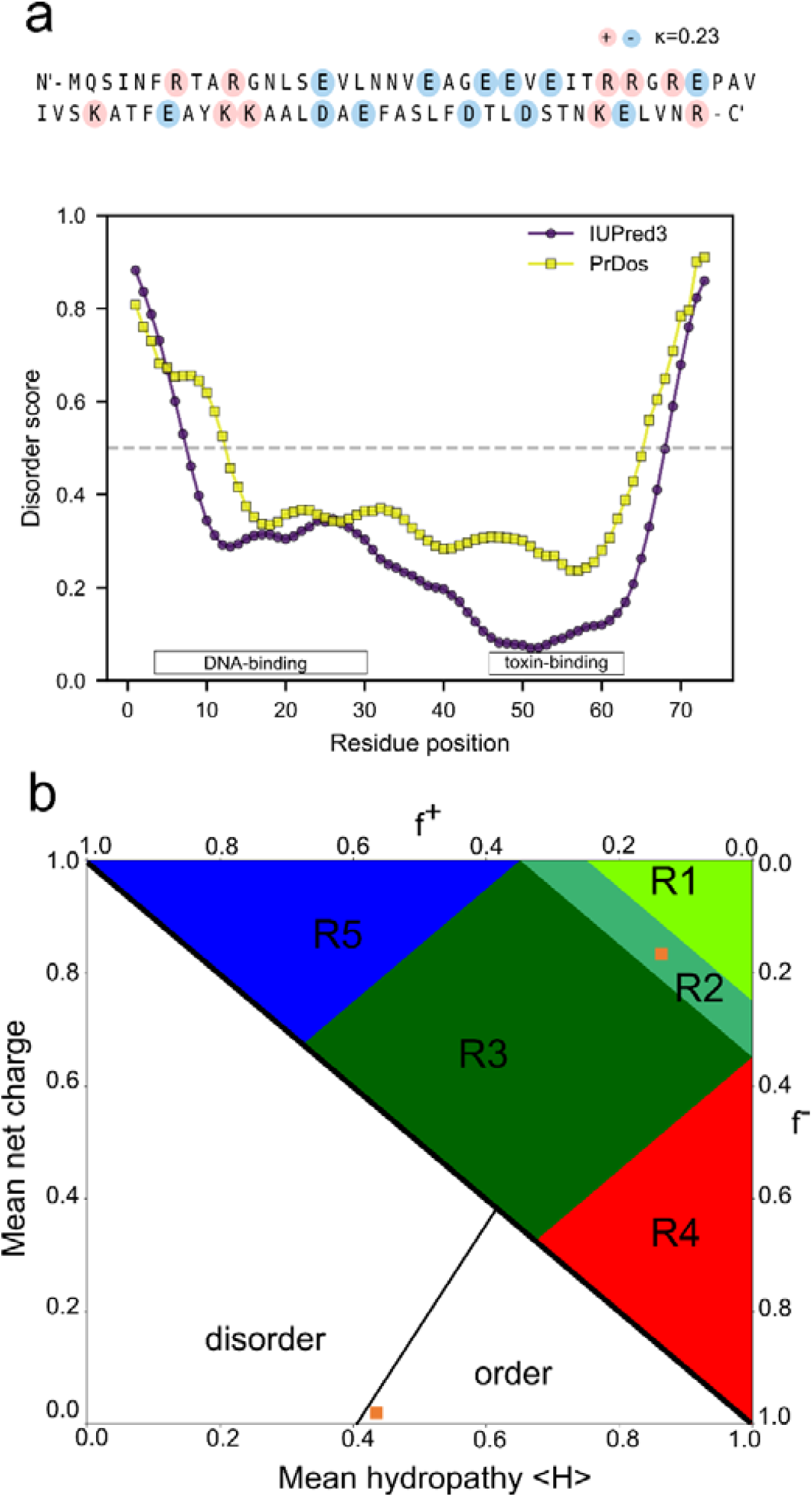
Phd transcription factor has “marginal” sequence properties. **a.** Distribution of charged residues in Phd. Positive and negative amino acids are colored in blue and red, respectively. Disorder prediction of Phd with the corresponding annotation toxin-binding and DNA-binding domains. **b.** Fraction of charged residues plot (upper half, right) and Uversky plot (lower half, left) show borderline properties of Phd (orange square), which is positioned between weak(R1)/strong(R3) polyampholytes and between disordered/ordered states.

The transcription regulator Phd from the *phd/doc* toxin-antitoxin (TA) module forms a good example to study such relationships as it combines disorder-to-order transition for both DNA recognition and neutralization of the toxin Doc, and both transitions are allosterically coupled. The *phd/doc* module was first discovered on bacteriophage P1, where it ensures the stable inheritance of the bacteriophage as a plasmid in the *E. coli* population (Lehnherr *et al*., 1993). The toxin Doc is a kinase that phosphorylates elongation factor Tu, thus stalling translation (Castro-Roa *et al*., 2013). Its activity is inhibited by the antitoxin Phd, with which it forms a complex (Magnuson & Yarmolinsky, 1998). This involves the C-terminal domain of Phd that is intrinsically unfolded in its free state and upon binding to Doc folds into a kinked α-helix that wraps around its partner (Garcia-Pino *et al*., 2008; Garcia-Pino *et al*., 2010). Phd is also a repressor that binds to the operator sequence of the *phd/doc* operon via its folded N-terminal domain (Magnuson & Yarmolinsky, 1998) This repression activity depends on the presence of Doc, with which it can form complexes of different stoichiometry (Garcia-Pino *et al*., 2010). In absence of Doc, the IDR regions of two adjacent Phd dimers on the operator interfere, resulting in weak binding (Garcia-Pino *et al*., 2016). At Doc:Phd ratios close to one, repression is strong, while at high Doc:Phd ratios, repression is relieved. This mechanism of regulation has been termed “conditional co-operativity” and is observed in a number of otherwise unrelated TA systems (Overgaard *et al*., 2008; De Bruyn *et al*., 2021).

Here, we show that Phd behaves as a prototypical “marginal” IDP, with sequence features that place it between disordered and ordered, as well as between collapsed and expanded IDPs. We demonstrate that Phd samples a diverse pool of disordered and partially ordered states whose populations depend on the salt concentration and the monomer–dimer equilibrium. Under near-physiological conditions, Phd resides close to a triple point in the phase diagram, where three macroscopically distinct ensembles coexist. Combined with previous crystallographic data, we chart the complete conformational space of Phd, spanning from fully disordered to fully ordered states. Our results highlight how marginal sequence properties of Phd link to its structural heterogeneity, which enables a range of functional disorder-to-order interactions with its partners essentially functioning as a conformational rheostat.

## RESULTS

### Sequence features place Phd antitoxin at the border between order and disorder

Recent advances in understanding the sequence-ensemble relationship of IDPs have shown that simple metrics, such as charge distribution and hydropathy, can capture global properties of IDP ensembles, including their compactness and the degree of order (Das & Pappu, 2013; Hofmann,*et al*., 2012). The Phd antitoxin is a 73-amino-acid-long protein with the DNA-binding domain at the N-terminus followed by the toxin-binding sequence (**Figure 1a**). The disorder prediction however, does not map to its domain arrangement; N and C termini are predicted as disordered while the central region is ordered (**Figure 1a**). Interestingly, AlphaFold3 predicts a fully ordered structure, with the N-terminal domain adopting a long α-helix (**Figure S1**), a conformation otherwise observed in the complex with Doc protein (Garcia-Pino *et al*., 2010). Approximately one third of the residues of Phd are charged, with a nearly equal proportion of positive and negative charges (10 positively and 12 negatively charged residues, **Figure 1a**). These charges show a relatively even distribution (mixing parameter κ=0.231, Das & Pappu, 2013) spanning both globular and disordered protein regions. The high fraction of charged residues, combined with comparatively low hydrophobicity, positions Phd near the empirical boundary between folded and disordered proteins on the Uversky plot (**Figure 1b**). Other metrics relevant to polyampholyte behavior, such as fraction of charged residues (FCR = 0.3) and the balance of charges, indicate that Phd behaves as an intermediate, neutral polyampholyte IDP (region R2 on **Figure 1b**). Consistently, it also falls within the boundary region on the Das-Pappu diagram, between collapsed and expanded conformations (**Figure 1b**). Taken together, these sequence characteristics suggest that Phd is a “marginal” IDP - borderline both in terms of order versus disorder and collapse versus expansion. This implies that its conformational ensemble is likely to be highly sensitive to changes in solution conditions.

### The conformational ensemble of Phd is sensitive to changes in ionic strength

We next examined how changes in ionic strength affect the conformational ensemble of Phd using circular dichroism (CD) spectroscopy and small-angle X-ray scattering (SAXS). Far-UV CD spectra of Phd in absence of added salt (buffer only) display typical features of a predominantly disordered protein (**Figure 2a**). Spectra show a weak minimum at 222 nm suggesting nascent α-helical structure and stronger minimum at low wavelengths typical for coiled conformation. Strikingly, progressive addition of NaCl produces dramatic changes in the spectra (**Figure 2a**), consistent with protein ordering and accompanying increase in α-helical content. The mean molar ellipticity at 222 nm, which correlates with the α-helicity, decreases from -2800°cm² dmol^-1^ in buffer to -5100 °cm^2^ dmol^-1^ in 500 mM NaCl (**Figure 2b**). Secondary structure estimation using the Bestsel algorithm (Micsonai *et al*., 2025) indicates that salt increases α-helical content from 2 to 12%, primary at the expense of turn structures (**Table S1**). We recently showed that helical content estimated from CD is underestimated when summation of CD signal neglects the ensemble nature of the proteins, which is particularly apparent in case of IDPs with only a small helix content (Zavrtanik *et al*., 2024). Using the ChiraKit server (Burastero *et al*., 2025) that applies ensemble-based CD signal summation for helix content estimation, and we observe an increase from 13% to 20% helicity due to increasing ionic strength (**Table S1**). To explore the mechanism behind salt-induced structuring of Phd, we repeated titrations using salts with different alkali metal cations (Li^+^, Na^+^, K^+^, Rb^+^, Cs^+^). If structuring were driven primarily by the hydrophobic collapse, cation-specific effects consistent with the Hofmeister series should be apparent, reflecting the involvement of hydrophobic residues and indirect ion effects on water properties (Brini *et al*., 2017). However, we observe practically identical behavior for all monovalent cations, indicating that the dominant effect is electrostatic, arising from charge screening, rather than modulation of hydrophobic effect (**Figure 2b**).

**Figure 2.**
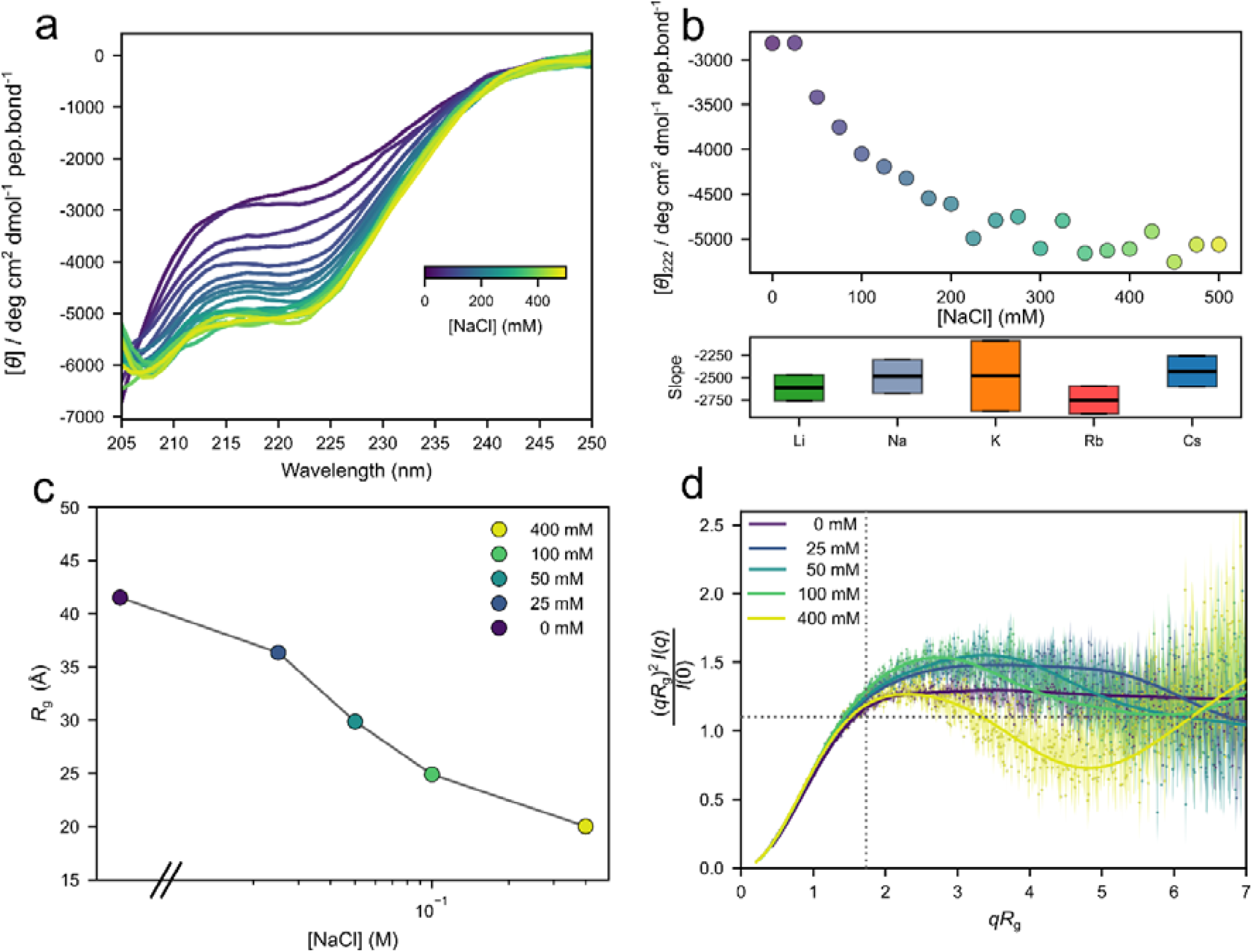
Salt promotes collapse and structuring of the disordered Phd ensemble. **a.** Circular dichroism (CD) spectra of Phd with increasing concertation of NaCl (from 0 to 500 mM) shows formation of secondary structure with increasing NaCl. **b.** The corresponding change in the CD signal intensity at 222 nm for NaCl titration, which correlates with amount of helix structure. Bottom panels compares the observed slopes (CD signal vs. log salt concertation) for titrations using different monovalent salts with chloride anion. **c.** Radius of gyration derived from Porod analysis of SAXS profiles (*R*_G_) at different NaCl concentrations. **d.** Dimensionless Kratky plots for Phd at different concentrations of NaCl indicates reduction of conformational flexibility with increasing NaCl concentration. The intercept at (√3, 1.104) marks the position of the maximum for an ideal globular protein.

We next used small-angle X-ray Scattering (SAXS) to delineate the effects of salt on the overall shape of the Phd ensemble. Increasing ionic strength induces collapse of the protein ensemble with the radius of gyration (*R*_G_) changing from 41.5 ± 0.4 Å in buffer only to 20.0 ± 0.2 Å in 400 mM NaCl (**Figure 2c**). Dimensionless Kratky plots show the accompanying changes in Phd flexibility as a function of salt (**Figure 2d**), with flexible chains in buffer only changing into more globular ensembles at 400 mM NaCl, as represented by the bell-shaped curve in the corresponding Kratky plot. Collectively, both CD and SAXS show that increase of ionic strength leads to the collapse of Phd ensemble to a more compact, structured one, mainly through to screening of electrostatic charges on the protein.

### Ionic strength modulates the Phd monomer-dimer equilibrium

Intrigued by the strong sensitivity of the Phd conformational ensemble to ionic strength, we next investigated which protein regions participate in the observed structural transitions. Previous crystallographic studies identified different dimeric states of Phd with varying degrees of disorder in the C-terminal helix (Garcia-Pino *et al*., 2010). However, the CD spectra at low salt concentrations (**Figure 2a**) reveal very little secondary structure - less than what would be expected for the Phd dimers observed cryptographically. Furthermore, salt-induced changes in the structural ensemble observed by CD and SAXS significantly exceed the modes variation of the C-terminal helix structure (Garcia-Pino *et al*., 2010).

We therefore hypothesized that at low ionic strengths and protein concentrations a transition into less structured, monomeric state is favored. To test for the presence of monomer-dimer equilibrium, we measured CD spectra at varying Phd concentrations and a fixed low ionic strength (buffer only, 0 M NaCl) (**Figure 3a**). After normalization by protein concentration, we observe changes in the CD spectra that are consistent with secondary structure formation coupled to a non-monomolecular process, most likely a monomer-dimer equilibrium. Increasing protein concentration leads to increased protein ordering, evidenced by a mean molar ellipticity at 222 nm decreasing from -3000° cm^2^ dmol^-1^ at 2 μM Phd to -7300° cm^2^ dmol^-1^ at 150 μM Phd. The changes in the CD signal intensity with protein concentration can be described using a dimer dissociation model, which yield an apparent dissociation constant of about 600 μM. We next repeated the Phd concentrations series in the presence of 400 mM NaCl (**Figure 3b**). As expected from the salt titration experiments (**Figure 2**), the starting state is more structured (MRE ≈ –4800 °cm² dmol⁻¹). Furthermore, in the presence of salt, the transition to a more structured ensemble occurs at lower Phd concentrations, consistent with a reduced apparent dissociation constant of about 200 μM. However, full saturation of the CD signal at maximal Phd concentrations was not achieved under the conditions tested, and further increase of protein concentration was not possible due to excess sample absorbance. Overall, these results indicate that free Phd exists in an equilibrium between an unstructured monomer and a more structured dimer, and their relative populations are modulated by the ionic strength.

**Figure 3.**
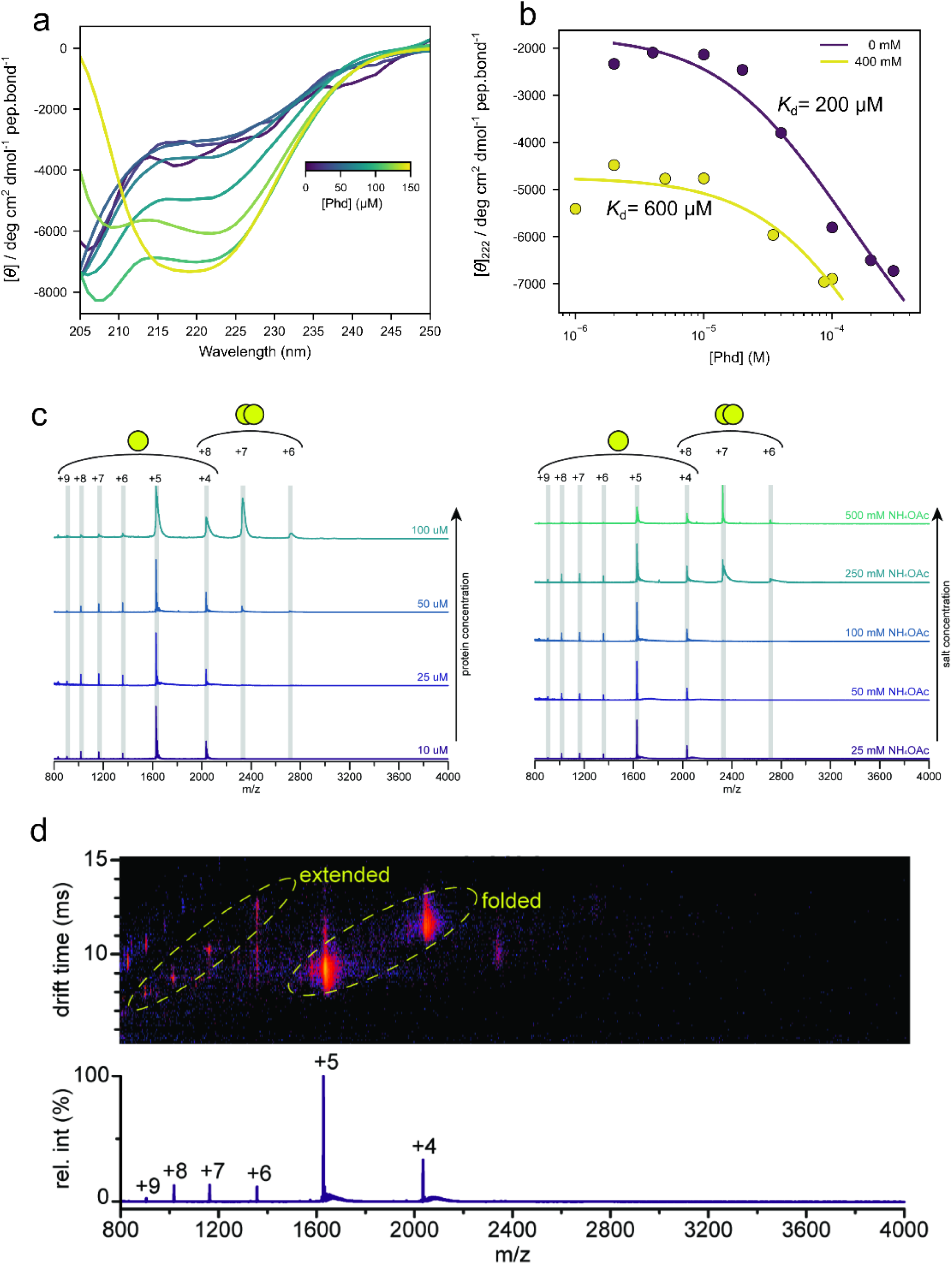
An equilibrium between monomeric and dimeric ensembles is modulated by salt. **a.** Normalized Phd CD spectra at varying concentration of protein indicate protein structuring upon increase in protein concentration. **b.** Changes in the CD signal intensity at 222 nm as a function of Phd concertation in absence (empty symbols) and presence of 400 mM NaCl (full symbols). Solid liens show the fits to the dimer dissociation model and the corresponding apparent dissociation constant. **c.** Native mass spectra of Phd at different protein (left) or salt concentrations (right). Charge distributions corresponding to monomers and dimers are indicated on top of the stacked spectra. d. Mass spectrum of Phd at 25 mM NH4OAc (bottom) including the corresponding drift plot (top). The drift plot shows two distinct populations for monomeric Phd, indicating the co-existence of a folded (+4/+5) and extended (+6 - +9) structure.

### Native ion mobility-mass spectrometry reveals an equilibrium between dimeric and compact/extended monomeric conformations

To gain further insight into the monomer-dimer equilibrium, we applied native ion mobility-mass spectrometry. Again, we first investigated dependence on protein concentration, by measuring various Phd concentrations (10, 25, 50, and 100 μM) at a fixed salt concentration (100 mM NH₄OAc). Low protein concentrations (10-25 μM) show monomeric Phd only, whereas dimeric conformations progressively increase at higher Phd concentrations (>50 μM) (**Figure 3c, left panel**). Monomeric Phd displays a non-typical Gaussian charge state distribution suggesting the coexistence of two structural conformations (**Figure 3c**). To investigate how salt modulates this tripartite equilibrium (compact/extended monomers and dimers), we acquired spectra at a constant protein (10 μM) but varying salt concentration (25-500 mM NH_4_OAc). Similar to what was observed at varying protein concentrations, a low ionic strength (25–100 mM NH₄OAc) results in compact and extended monomers only (**Figure 3c, right panel**) and the prevalence of dimeric conformations increase with the salt concentration. Moreover, extended/disordered monomers almost entirely disappear at 500 mM NH₄OAc, indicating a stabilizing effect of high ionic strength on the monomeric structure. This indicates that salt promotes the formation of compact monomeric and dimeric conformations, which agrees with results from CD spectroscopy. To confirm the coexistence of compact and extended monomeric species, we investigated their ion mobility profile (**Figure 3d**), which depends on mass, charge and shape, reflected by the rotationally averaged collision cross-section (CCS, Lanucara *et al*., 2014; Smith *et al*., 2009). CCS calculations revealed marked differences among monomer charge states, ranging from 1219 Å² (+5 monomer) to 1807 Å² (+7 monomer). The CCS of the monomeric +5 state is in close proximity to ubiquitin, a protein of similar size to Phd (May *et al*., 2018), indicating that this monomeric form of Phd is relatively compact. Altogether, this indicates the coexistence of a compact and more extended monomeric protein conformation of Phd.

### Thermodynamic analysis of the three-state equilibrium modulated by ionic strength

The data above suggest a model in which increase in ionic strength promotes the collapse of disordered monomers into a more compact, structured monomeric state, which then facilitates formation of Phd dimers. To disentangle the linkage between ionic strength, conformational ensemble and oligomeric state, we analyzed the thermal stability of Phd as a function of salt concentration using CD spectroscopy. Upon addition of salt the absolute CD signal intensity increases, but we also observe a strong increase in protein thermal stability (**Figure 4a**). The melting temperature increases from 23 °C in the absence of added salt (buffer only) to 56 °C in the presence of 0.8 M NaCl (**Figure 4a**). A complete recovery of the CD signal after renaturation indicates a reversible unfolding transition (**Figure S2**). Thermal denaturation of Phd was also followed by differential scanning calorimetry (DSC) at higher protein concentration (90 μM; **Figure 4b**), where reversible transitions were observed only in 200 mM NaCl, but not in absence of salt (**Figure S2**).

**Figure 4.**
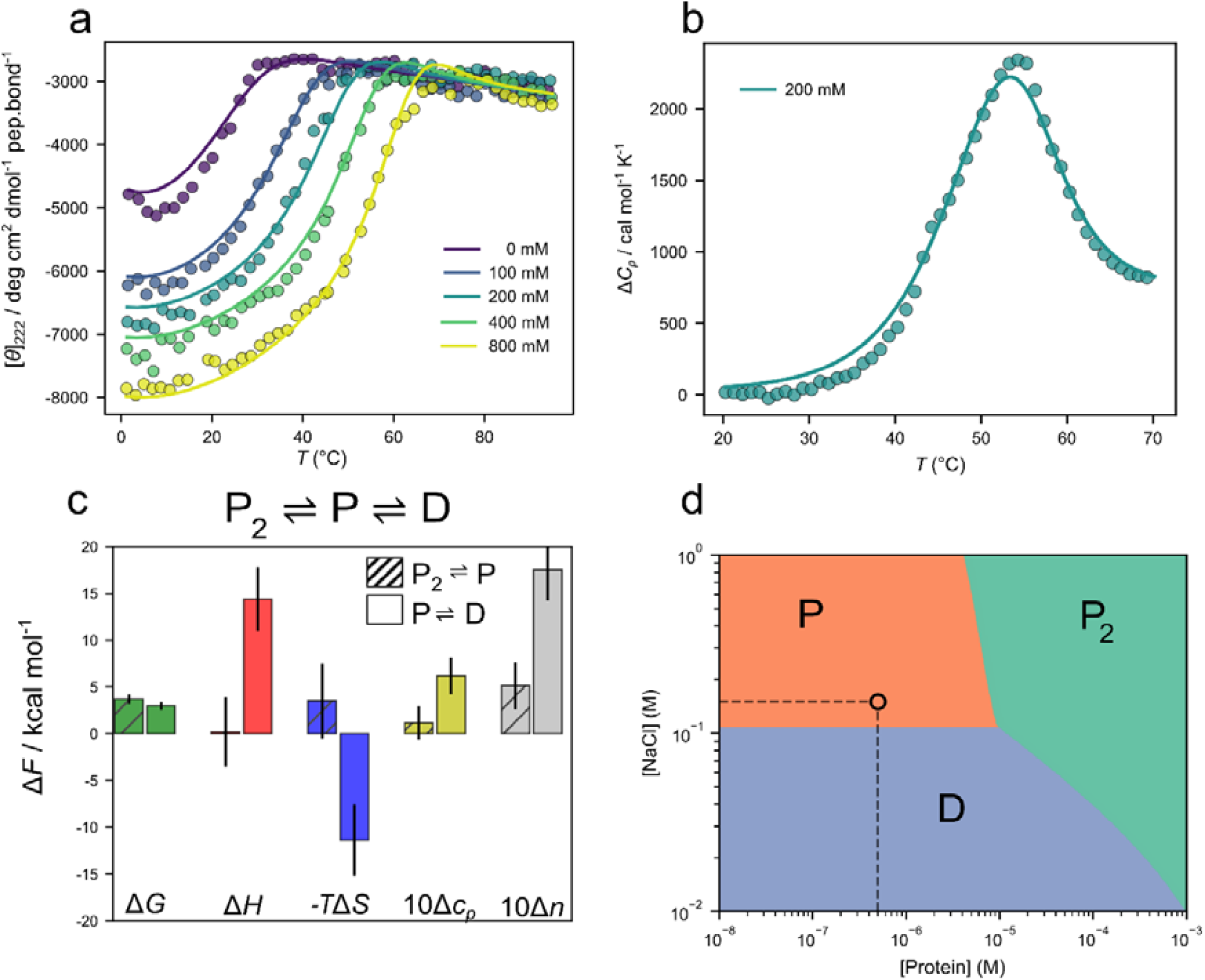
Phase diagram and thermodynamic stability Phd as a function of salt. **a.** Thermal melts of Phd monitored by CD show an increase in thermal stability with NaCl concertation. Solid lines represent the of the global fit of the three-state model to the experimental data. **b.** DSC scan of Phd in 20 mM Tris pH 7.5 and 200 mM NaCl. Global fit of the three-state model is shown as solid line. **c.** Three-state thermodynamic models assumes an interconversion of Phd dimer (P_2_) into partially structured monomer (P) and then to the disordered monomer (D). Bars compare the thermodynamic parameters for the two transitions obtained from the global fitting of the three state models to the CD and DSC data, with lines showing 1 std. Heat capacity change is reported in kcal mol^-1^ K^-1^ and Δ*n* representing number of released ions. **d.** Phase diagram of Phd shows the dominant macroscopic ensemble as a function of protein and salt concertation at 37 °C based on the parameters from the three-state thermodynamic model. Dashed lines show an approximate position corresponding to physiological solution conditions.

Thermal melts were described using a three-state model in which Phd exists in equilibrium between an unfolded monomer (M), a partially folded monomer (P) and a dimer (P_2_) (**Figure 4c**). The temperature dependence of the two equilibrium constants was described through the corresponding enthalpy and heat capacity change, while the salt dependence was modeled using an ion-exchange framework, where the free energy depends logarithmically on ion activity scaled by the number of bound ions (see *Methods*) (Tan & Chen, 2006; Vancraenenbroeck *et al*., 2019). The three-state model provides a satisfactory description of the experimental data and yields reasonable parameter values (**Figure 4c, Table S2**). At standard conditions, dissociation of dimers into partially structured monomers is practically isoenthalpic (Δ*H*=0.2 kcal/mol) but entropically unfavorable (-*T*Δ*S*= +3.5 kcal/mol), which is unexpected. On the other hand, transition from the partially structured monomer into unfolded monomer has a thermodynamic signature that would be expected for protein denaturation: an unfavorable enthalpy Δ*H*=+14.4 kcal/mol, favorable entropy change -*T*Δ*S*= - 11.4 kcal/mol and a positive change in heat capacity Δ*c_P_=*0.6 kcal/molK). However, the absolute values are smaller compared to an average globular protein (Robertson & Murphy, 1997), supporting the molten-globule character of this state. Denaturation of the partially folded monomer is further accompanied by the release of 2 ions per monomer (**Figure 4c**). Thus, increasing salt concentration is associated with binding of two ions and leads to collapse of monomers to what is likely a molten globule form.

Based on these thermodynamic parameters, we determined the most probable Phd state at given solution conditions (temperature, protein and salt concentration). This is represented in Phd phase diagram (**Figure 4d**), where the dominant conformations are represented as colored regions. The border lines indicate coexistence between two states, while the triple point represents solution condition where all three states are present at the same molar fraction. Inspection of the salt versus protein concentration diagram shows that at 37 °C and the physiological salt concentrations (150 mM NaCl), Phd exists predominantly as a monomer up to ∼5 μM protein concentration. Change in the concentration of salt modulates the conformation of monomers between disordered and compact forms (vertical movement on **Figure 4d**). At micromolar protein concentrations and above the dimer state dominates, which can however dissociate into monomers when salt concentration is decreased (**Figure 4d**).

### SAXS-derived ensembles reveal a gradient of disorder in Phd dimers

To investigate the structural basis of salt modulation, we attempted to model the conformational ensemble of Phd using experimental SAXS data. As a starting point, we used the crystal structure of the Phd dimer, which is fully ordered in complex with the Doc toxin (PDB: 3KH2). In this structure, the majority of the dimer interface is formed through the DNA-binding domain, while the C-terminal toxin-binding domain adopts an extended helix followed by a short-kinked helix at the C-terminus. However, the calculated SAXS profile based on this crystal structure did not fit the experimental scattering curves obtained at high salt concentration (400 mM NaCl), which we reasoned is due to the presence of residual Phd monomers. We next estimated monomer and dimer fractions at conditions under the protein and salt conditions used for SAXS, using the parameters obtained from the three-state thermodynamic model. However, even now the fraction-weighted theoretical scattering profiles failed to reproduce the experimental SAXS curves, which consistently suggested an ensemble that was overall more extended than predicted.

We next hypothesized that additional disorder may arise from Phd dimers themselves, which might have more disorder than our starting dimer model taken from the Phd-Doc crystal structure. To test this, we systematically introduced disorder into Phd dimers by progressively allowing full flexibility in 10-residue segments starting from the C-terminus, while keeping the remainder of the protein fixed. This procedure generated six dimer variants with different degrees of disorder. We then re-calculated SAXS curves for ensembles consisting of the fraction-weighted sum of monomers and dimers, but using each time a different dimer variant. We then selected the appropriate dimer variant in the ensemble based on the lowest χ² value (**Figure S3**). At the highest salt concentrations, the ensemble contains a small fraction of monomers (∼12%), while the dimers are largely ordered with only the last 10 residues exhibiting flexibility (**Figure 5a**). As the ionic strength decreases, the monomer fraction increases to ∼30%, and the C-terminal helical domain of the dimers became progressively more disordered (**Figure 5b-d**). Thus, in addition to the structural transitions associated with the monomer-dimer equilibrium identified by CD spectroscopy and Native MS-Ion Mobility spectroscopy, SAXS-based modeling indicates further gradation of the disorder in the dimer state itself.

**Figure 5.**
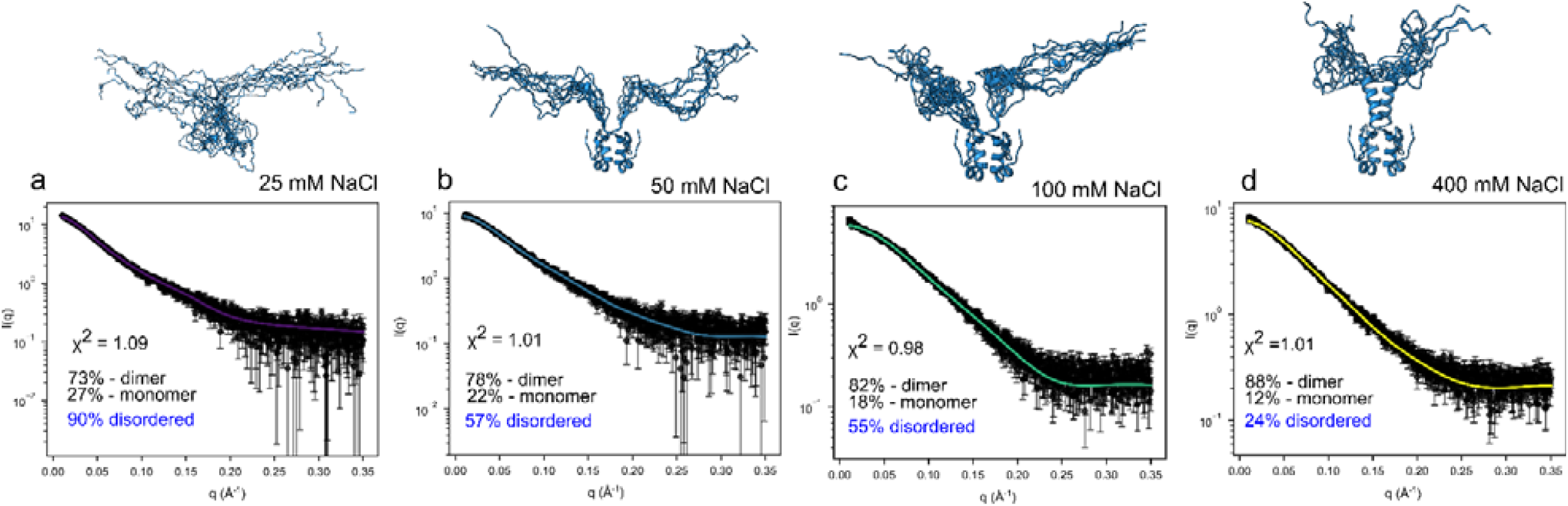
SAXS-derived ensemble of Phd dimer shows gradation of disorder. **a.-d.** Experimental SAXS data (black dots) and back-calculated curves (colored line) for the best refinement solutions at a given monomer/dimer ratio (derived from the thermodynamic model) and different salt concentrations. The χ2 values for the best fits, the relative populations of the Phd monomers and dimers, as well as the total amount of the disordered residues in the system are indicated in the plots. The insets show ensembles of 10 best solutions for the Phd dimer, with the increasing degree of structural disorder inversely proportional to the solution ionic strength.

## DISCUSSION

The Phd antitoxin is a prototypical disordered transcription factor, in which structural disorder plays a key function in the allosteric transcriptional regulation of the toxin–antitoxin module (Garcia-Pino *et al*., 2010; Garcia-Pino *et al*., 2016). Here, we reveal the presence of additional monomeric and dimeric disordered states at physiological conditions of Phd. Together with previous crystallographic data, these findings allow us to comprehensively chart the structural space of Phd (**Figure 6**), spanning from fully disordered to completely ordered states. To our knowledge, this represents one of the most detailed catalogues of a disorder gradient within a single protein. The degree of disorder is closely coupled to the oligomeric state of the protein, showing that homo- and heteromeric interactions progressively promote structural ordering in Phd (**Figure 6**). External factors such as temperature and co-solutes like salt modulate this balance, thereby extending the structural repertoire of Phd and enabling mapping of its conformational states.

**Figure 6.**
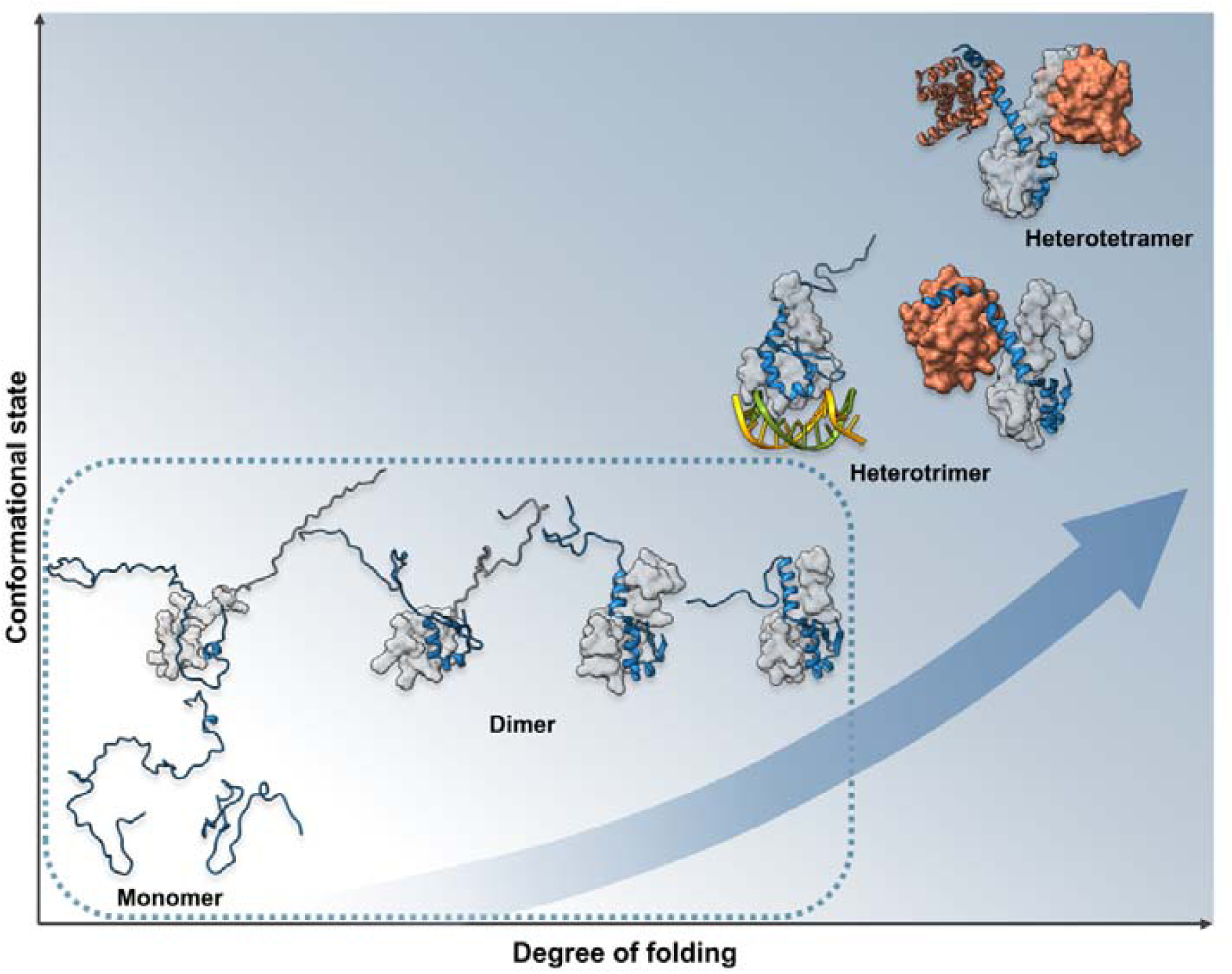
Cartography of Phd structural ensemble shows an order-disorder continuum. Snapshots from the Phd structural ensemble as a function of oligomerization state and the degree of structuring. For clarity only a single Phd molecule is shown as cartoon, partner molecule (second Phd molecule or Doc toxin) is shown as surface. Encircled are states that correspond to the pool of free Phd conformations available for functional interactions with partner molecules. Monomeric states are shown schematically based on the results from CD and native mass spectroscopy. Dimeric states correspond to SAXS data at low salt concentration, followed by the last two dimers observed crystallographically (PDB 3HS2 and 3HRY). More structured higher-order assemblies with Doc toxin and DNA operator sequence correspond to crystallographic models 4ZM0, 3K33 and 3KH2.

At low protein concentrations in absence of salt Phd exists as disordered, extended ensemble, as evidenced by CD spectra, large collision cross-section from native mass ion mobility spectroscopy, large radius of gyration and high degree of disorder evidenced by Kratky plots (**Figure 2, 3**). Collapse into a more compact monomeric state is observed by increasing ionic strength, likely through electrostatic screening, with lack of ion-specificity (**Figure 2b**). In contrast to cases where binding of disordered proteins results in entropically favorable release of salt ions (Chowdhury *et al*., 2023), monomer structuring and homo-dimerization are both accompanied by ion binding. This is more similar to a process such as polyelectrolyte folding (Manning, 1978). Ion mobility spectroscopy CCS values of this partially folded monomer are similar to that of a dimer, whereas its CD signal is only slightly more intense compared to the disordered monomer. Taken together these results indicate a compact monomeric state with very little secondary structure, likely a molten-globule type of structure. This is also supported by the thermodynamic signature of the partial monomer unfolding (**Figure 4c**), with the qualitative signature similar to protein unfolding, but overall diminished values relative to an average globular protein.

Increasing salt or protein concentration promotes homodimerization of Phd, which is accompanied by a gain in structural order. Previous crystallographic studies have identified several distinct dimeric conformations of Phd (Garcia-Pino *et al*., 2010). These conformations range from fully structured to only limited structure in the N-terminal domain. Our SAXS models as a function of ionic strength closely follow these degrees of structuring. For example, in a crystal form obtained at low ionic strength, one Phd molecule shows a loosely packed hydrophobic core and no well-defined secondary structure. Such a conformation closely corresponds to our SAXS model of Phd at low salt concentration (**Figure 5d**). Highly ordered conformations on the other hand are favored by high ionic strength and additional interactions with Doc and/or DNA. This structuring is allosteric: DNA binding induces structure in the C-terminal IDR while Doc binding further stabilizes the N-terminal domain. Taken together, these crystallographic snapshots, combined with our SAXS-based models, reveal a continuum of conformations in Phd characterized by a gradation of disorder that extends from the C-terminus towards the N-terminus (**Figure 6**).

Our findings suggest that functional interactions of Phd are directly coupled to its degree of disorder. Transcriptional regulation occurs through an allosteric mechanism, in which Doc binding induces structuring of not only the C-terminal IDR of Phd, but also its N-terminal DNA binding domain. This allosteric structuring is accompanied by an increase in DNA operator affinity (Garcia-Pino *et al*., 2010). The present study adds an additional layer of complexity by identifying three novel macroscopic ensembles of free Phd with different amounts of disorder. All together this represents a large and diverse pool of conformations available for Doc or DNA binding and extends the range of disorder-to-order transitions that can occur in Phd upon binding. The distribution of these ensembles is highly sensitive to solution conditions, as illustrated by the phase state diagrams in **Figure 5c**. Such sensitivity may allow dynamic redistribution of binding-competent states in response to environmental changes. Although the exact intracellular concentration of Phd is not known, protein abundance is typically on the order of a few hundred copies per cell roughly corresponding to micro-to-sub micromolar concentrations (Milo et al., 2016). Under physiological salt concentration, this is not far from the triple point in the Phd phase diagram (**Figure 4d**), where unfolded monomer, partially folded monomer, and dimer are populated in approximately equal proportions. Not only its amino acid sequence properties, but also the biophysical properties place Phd at the stability “triple point,” where it can readily interconvert between different conformational and oligomeric states. Such positioning would therefore enable Phd to function as a conformational rheostat: interconversion between its different structural ensembles coupled to monomer-dimer equilibrium enables more gradual ensemble shifts upon binding to Doc or operator DNA (Muñoz *et al*., 2016). A similar rheostatic behavior was observed in another transcription factor NCBD, which was reported to function as allosteric regulator through interconversion between different structural ensembles (Luong *et al*., 2022).

Overall, this study accurately maps the conformational plasticity of Phd transcription factor. We link this heterogeneity to the marginal sequence properties that position Phd protein at a threshold between order and disorder. It is possible that the molecular mechanism governing the rheostatic conformational behavior of Phd represents a general theme for the marginally disordered transcription factors.

## EXPERIMENTAL PROCEDURES

### Protein production and purification

Recombinant expression and purification of Phd was carried out essentially as described in Garcia-Pino *et al*. (2010) and Sterckx *et al*. (2015) with minor modifications. The pET21b plasmid harboring both Phd and Doc encodes a C-terminal His-tag on Doc, while Phd remains untagged. A single colony of *E. coli* BL21 (DE3) transformed with pET21b-*phd/doc* was inoculated into 120 ml LB medium supplemented with 100 µg/ml ampicillin, 1% glucose and grown overnight at 37°C with shaking. The overnight culture was diluted 100-fold in 12 l LB medium with ampicillin, 0.5% glucose and grown at 37°C with shaking. When OD_600_ of the culture reached 0.8-0.9, expression of the operon was induced with 1 mM IPTG. After further incubation for 4 h, the cells were harvested by centrifugation at 5000 rpm for 20 min at 4°C using a JLA-8.1000 rotor. The pellets were resuspended in lysis buffer (50 mM Tris, pH 8.0, 500 mM NaCl, 0.1 mg/ml AEBSF and 1 µg/ml leupeptin) and the suspension was then left to stir at 4°C for 30 min. The cells were lysed using a cell cracker and the lysate was cleared by centrifugation at 16000 rpm for 30 min at 4°C using a JA20 rotor. The supernatant was loaded onto a 5-ml HisTrap HP nickel-Sepharose column that had been pre-equilibrated with at least five column volumes of Buffer A (50 mM Tris, pH 8.0, 500 mM NaCl). After thorough washing of the column with Buffer B (50 mM Tris, pH 7.5 and 1 M NaCl, 10% ethylene glycol), Phd was dissociated from Doc by applying a step gradient (0.0, 2.5 and 5.0 M) of guanidine hydrochloride (GnHCl) in 50 mM Tris, pH 7.0 and 500 mM NaCl. The fractions containing Phd were pooled, diluted 10 times in 50 mM Tris, pH 7.5, 500 mM NaCl and dialyzed overnight against the same buffer to get GnHCl removed. For further purification, the protein was concentrated to a volume of 2 ml and subsequently loaded onto a Superdex 75 16/60 HR column that had been pre-equilibrated with SEC buffer (50 mM Tris, pH 7.5 and 200 mM NaCl). The eluted fractions were resolved by SDS PAGE and the relevant fractions were pooled, flash-frozen and stored at -80°C until further use.

### Circular dichroism spectroscopy

All CD measurements were performed on the JASCO J-1500 spectrometer. Thermal melts were recorded in a cuvette with 1 mm optical pathlength with 20 μM Phd solution in 20 mM Tris pH 7.5 and different concentration of NaCl (0, 80, 200, 400, 800 mM). CD melting curves were recorded between 4^◦^C and 96^◦^C by measuring the intensity at the 222 nm in intervals of 1^◦^C at a speed of 1^◦^C/min. All measured signals were converted to molar ellipticity. Titrations of salt into solution containg Phd were recorded in cuvette with 5 mm optical path length at 20 °C using salt stock solution 3 M. Dependence of the CD signal intensity on the Phd concertation was recorded at 20 °C using 5 mm (low concentration of Phd) or 1 mm (above 40 μM Phd concentrations) cuvette. CD spectra were analyzed using Bestsel (Micsonai *et al*., 2025) and Chirakit (Burastero *et al*., 2025) webservers using default parameters.

### Differential scanning calorimetry (DSC)

Differential scanning calorimetry (DSC) experiments were conducted using a Nano-DSC instrument (model 60200, Waters Ltd.) at a scanning rate of 1.4 K/min under elevated pressure of 3 atm. Prior to measurements, samples were degassed for 10 minutes and then loaded into the instrument’s cells. Raw data was converted to molar heat capacity using NanoAnalyze software, with setting partial specific volume (PSV) to 0.73 mL/g.

### Global fitting of three-state thermodynamic model

Mass-balance equations that describe interconversion of Phd between three macroscopic states were derived based on the model schematically shown on Figure 4C. Populations of disordered monomer, partially disordered monomer and a dimer can be expressed using two equilibrium constants and the solution is found analytically. Each equilibrium constant is related to the corresponding Gibbs free energy change, which depends on solution conditions:

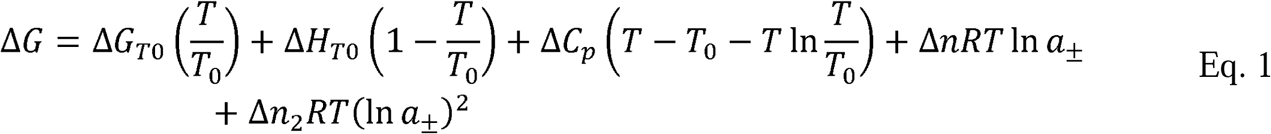

Here *T*_0_ is the reference temperature (*T*_0_=25 °C), while Δ*G*_T0_ and Δ*H*_T0_ are the standard free energy and enthalpy change at the reference temperature. Temperature dependence is given by standard thermodynamic expressions and assuming temperature-independent heat capacity change Δ*C_p_*. Salt dependence was described by the counterion release model with parameter Δ*n* corresponding to the number of exchanged ions (Record *et al*., 1978). To account for experiments performed at higher salt concentrations (above 200 mM), an additional quadratic term was included in Eq. 1 (Owczarzy *et al*., 2004). Furthermore, salt activity, rather than salt concentration, was used in data analysis. Activities have been calculated using experimental activity coefficients (Robinson & Stokes, 1970). The thermodynamic master equation (Eq.1) defines that equilibrium constant at given solution conditions, which are used to determine the population of species through mass-balance equations. These are directly related to the observed spectroscopic and calorimetric signals by:

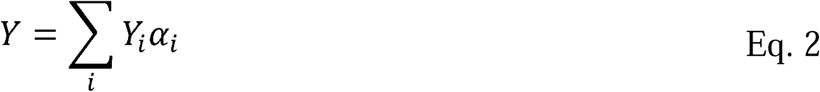

Here *Y* corresponds to the overall signal, which is given as sum of the *Y*_i_ (signal contribution of i-th species) weighted by the equilibrium fraction of i-th species α_i_. In case of CD spectroscopic signals *Y*_i_ are the mean molar ellipticities of species i (i = P_2_, P or D) which depend linearly on temperature. Temperature dependence was extracted from the post-transition baseline in case of species D or used as fitting parameter. In the case of DSC experiments *Y*_i_ correspond to the partial molar enthalpy of the sample, whose temperature-dependence is given by the corresponding heat capacity change.

### Small-angle X-ray Scattering

Synchrotron Radiation Small-angle X-ray scattering (SAXS) data from Phd in different salt concentrations were collected on the SWING beamline at the SOLEIL synchrotron (Gif-sur-Yvette, France) in batch mode. For each salt concentration, scattering was measured at three different protein concentrations (2-5 mg/ml) at 20°C. Prior to the measurement, samples were dialyzed against 20 mM Tris (pH 7.5), 2 mM EDTA and different concentrations of NaCl (0, 25, 50, 100, 150, 300, 400 mM). Following each measurement, the recorded frames were checked for radiation damage, and data from all concentrations were analyzed to exclude intermolecular events. Data were processed using the beamline software Foxtrot 3.5.2 (David & Perez, 2009) and analyzed with ATSAS (Manalastas-Cantos *et al*., 2021).

All simulations were performed in Xplor-NIH v 2.49 (Schwieters *et al*., 2003; Schwieters *et al*., 2006), starting from the crystal structure of the PhD dimer (PDB 3KH2). Missing (hydrogen) atoms were added in Xplor-NIH, followed by minimization of the energy function comprising the standard geometric (bonds, angles, dihedrals, and impropers) and steric (van der Waals) terms. To generate a starting monomer structure, a single chain of the PhD dimer was retained. An ensemble refinement of the PhD monomers and dimers, weighed by their respective relative populations (calculated form the mass-balance equations using parameters reported in Table S2), was performed against the experimental SAXS data following a published protocol (Deshmukh *et al*., 2013).

At each salt concentration, multiple refinement runs were performed, where the PhD monomer was treated as fully flexible (by allowing full translational and rotational degrees of freedom), while the PhD dimer was progressively unfolded in steps of 10 residues starting from its C-terminus. In particular, six refinement runs were carried out each time, where PhD dimer residues 63-73, 53-73, 41-73, 33-73, 23-73, 12-72 were given the full degree of freedom, while the rest of the protein was kept fixed (these correspond to 14, 28, 45, 56, 70, and 86 % of PhD dimer unfolded). The computational protocol comprised an initial simulated annealing step followed by the side-chain energy minimization as described before (Schwieters & Clore, 2014). In addition to the standard geometric and steric terms, the energy function included a knowledge-based dihedral angle potential and the SAXS energy term incorporating the experimental data (Schwieters *et al*., 2018). Truncated SAXS curves (q < 0.35 Å^-1^) were used as the sole experimental input.

In each refinement run, 100 structures were calculated and 10 lowest-energy solutions – representing the best agreement with the experimental data – retained for the subsequent analysis. The agreement between the experimental and calculated SAXS curves (obtained with the calcSAXS helper program, which is part of the Xplor-NIH package) was assessed by calculating the χ^2^:

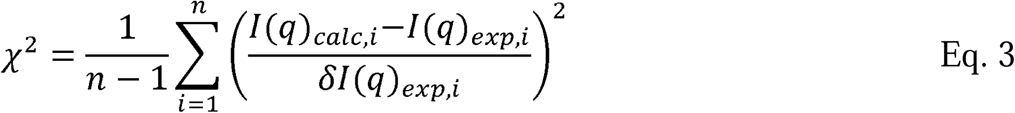

where l(q)_calc,i_ and l(q)_exp,i_ are the scattering intensities at a given q for the calculated and experimental SAXS curves, ol(q)_exp,i_ is an experimental error on the corresponding value, and n is the number of data points defining the experimental SAXS curve. Comparison of the χ^2^ values for the best solutions at different runs simulating varying degrees of PhD dimer’s structural disorder allowed to define the minimal amount of the dimer unfolding that is required to obtain good agreement (low χ^2^) with the experimental SAXS curves at different salt concentrations.

### Native MS and ion mobility analysis

Native MS experiments require volatile buffer salts to prevent adverse effects such as ion suppression and/or adduct formation (Konijnenberg *et al*., 2013). Thus, Phd was buffer-exchanged to a volatile ammonium acetate (NH_4_OAc) solution using P-6 micro Bio-Spin columns (Bio-Rad). To study the effect of both protein and salt concentration, Phd was prepared at varying protein as well as NH_4_OAc concentrations. The protein concentration was ranged from 10-100 µM (10, 25, 50, and 100 µM) in 100 mM NH_4_OAc and the salt concentration was ranged from 25-500 mM NH_4_OAc (25, 50, 100, 250, 500 mM) using a Phd concentration of 10 µM. For native ion mobility-mass spectrometry (IM-MS) experiments, a few microliters of each sample were loaded into in-house produced gold-coated borosilicate glass capillaries to facilitate infusion into a Synapt G2 HDMS (Waters, Wilmslow, UK) mass spectrometer. Nano-electrospray ionization (nanoESI) was applied to generate protein ions, which were drawn into the vacuum of the mass spectrometer. The instrument was operated in mobility mode and the following crucial parameter settings were applied to retain the native, solution-phase structure of the protein: 1.4 kV capillary voltage, 25 V sampling cone, 1 V extractor cone, 15 V and 2 V collision energy in the trap and transfer collision cell, respectively, and 45 V trap DC bias. IM parameters were set to 600 m/s wave velocity, 35 V wave height, and a gas flow of 180 mL/min and 90 mL/min in the helium and ion mobility cell, respectively. Pressures throughout the instrument were maintained at 4.7 mbar for backing, 3.0 mbar (N_2_) in the ion mobility cell, and 0.027 mbar (Ar) in the trap and transfer collision cell. Collision cross sections (CCS) were calculated after calibrating ion mobility arrival times by means of native measurements of the following standard proteins: cytochrome c (monomer: +6), β-lactoglobulin (monomer: +7/+8, dimer: +11/+12), avidin (tetramer: +15/+16/+17), and bovine serum albumin (monomer: +14/+15/+16).

