## Supplementary material for "Ionic strength modulates structural disorder and protein oligomerization in the marginally disordered Phd transcription factor"

for

**Table S1**. Deconvoution of CD spectra using Bestsel algorithm for determination of secondary structures of Phd at different protein concentrations in 20 mM Tris pH 7.5 (the corresponding CD spectra are shown on Figure 2A), or for 10 µM Phd concentration at different NaCl concentrations.

| protein concentration (µM) | **2** | **4** | **10** | **20** | **40** | **100** | **200** | **300** |
| --- | --- | --- | --- | --- | --- | --- | --- | --- |
| **helix** | 0.031 | 0.035 | 0.036 | 0.083 | 0.083 | 0.079 | 0.119 | 0.144 |
| **antiP** | 0.268 | 0.297 | 0.203 | 0.264 | 0.237 | 0.319 | 0.311 | 0.279 |
| **P** | 0 | 0 | 0 | 0 | 0 | 0 | 0 | 0.003 |
| **Turn** | 0.17 | 0.173 | 0.199 | 0.075 | 0.182 | 0.162 | 0.151 | 0.154 |
| **O** | 0.531 | 0.495 | 0.562 | 0.478 | 0.497 | 0.44 | 0.419 | 0.421 |

| salt concentration (mM) | **0** | **136** | **275** | **388** | **500** |
| --- | --- | --- | --- | --- | --- |
| **helix** | 0.024 | 0.078 | 0.11 | 0.114 | 0.119 |
| **antiP** | 0.278 | 0.29 | 0.303 | 0.297 | 0.293 |
| **P** | 0 | 0 | 0 | 0 | 0 |
| **Turn** | 0.182 | 0.162 | 0.155 | 0.155 | 0.156 |
| **O** | 0.516 | 0.47 | 0.432 | 0.435 | 0.433 |

**Table S2.** Best-fit thermodynamics parameter for transition steps P to D (index 1) and P_2_ to P (index 2) at standard state conditions (I=1 M NaCl and *T*_0_=25 °C) given per 1 mol of monomer, as shown in Figure 4c. Here, Δ*G* and Δ*H* represent free energy and enthalpy and Δ*Cp* corresponds to the heat capacity change. Parameters g1 and g2 represent mean molar ellipticity at 222 nm for the pure P and P_2_ states, respectively. Parameters g1' and g2' represent their temperature-dependence.

| Δ*G*_1_ | (3000 ± 460) cal mol^-1^ | *ΔG*_2_ | (3700 ± 500) cal mol^-1^ |
| --- | --- | --- | --- |
| Δ*H*_1_ | (14400 ± 3500) cal mol^-1^ | Δ*H*_2_ | (200 ± 3700) cal mol^-1^ |
| Δ*C_p_*_,1_ | (620 ± 200) cal mol^-1^K^-1^ | Δ*C_p_*_,2_ | (120 ± 190) cal mol^-1^K^-1^ |
| Δ*n*_1_ | 2.1±0.3 | Δ*n*_2_ | 0.1±0.3 |
| Δ*n*_1'_ | 0.21 ± 0.04 | Δ*n*_2'_ | 0.05±0.04 |
| g1 | (-3900 ± 920) deg dmol^-1^ pep.bond^-1^ | g2 | (-10000 ± 1900) deg dmol^-1^ pep.bond^-1^ |
| g1' | (60 ± 30) deg dmol^-1^ pep.bond^-1^ °C^-1^ | g2' | (40 ± 50) deg dmol^-1^ pep.bond^-1^ °C^-1^ |

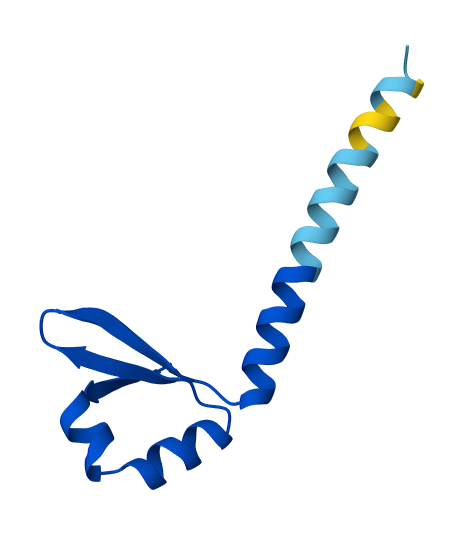

**Figure S1.** Three-dimensional structure of Phd as predicted by AlphaFold 3 (AF3).

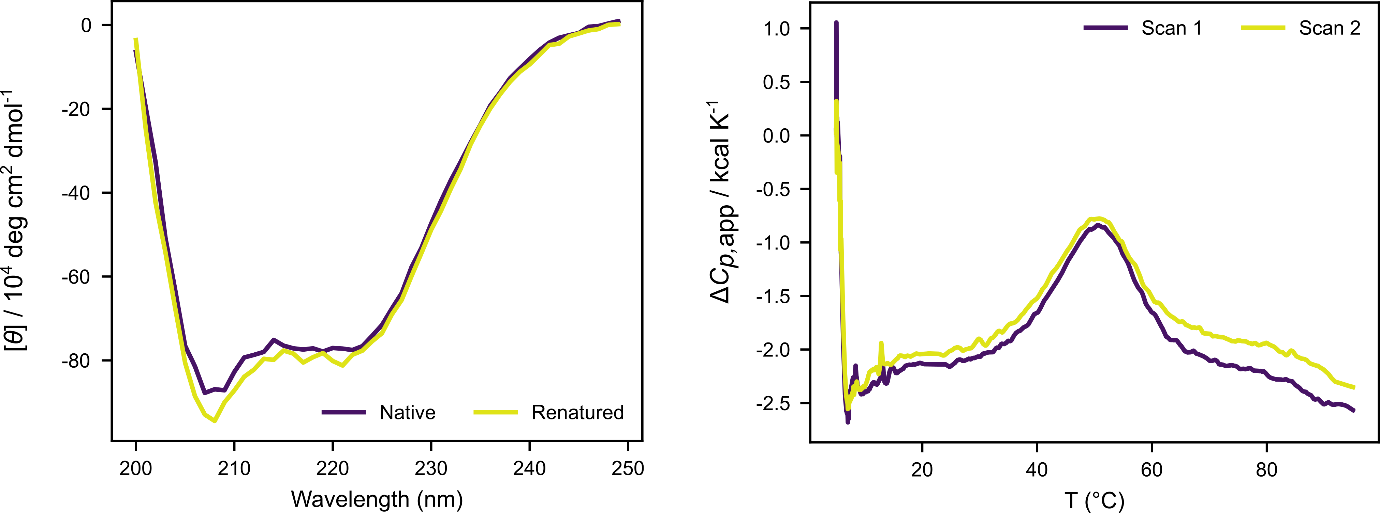
**Figure S2. Reversibility of Phd denaturation by CD spectroscopy and DSC.**

Two CD spectra: native recorded before the thermal denaturation experiment and the other (renaturated) recorded after the experiment, show that transition is reversible. Two consecutive DSC scans (apparent heat capacity scale) of Phd in 200 mM NaCl and 86 µM concentration are very similar, suggesting a reversible process. On the other hand DSC scans at lower salt concentrations are not reversible.

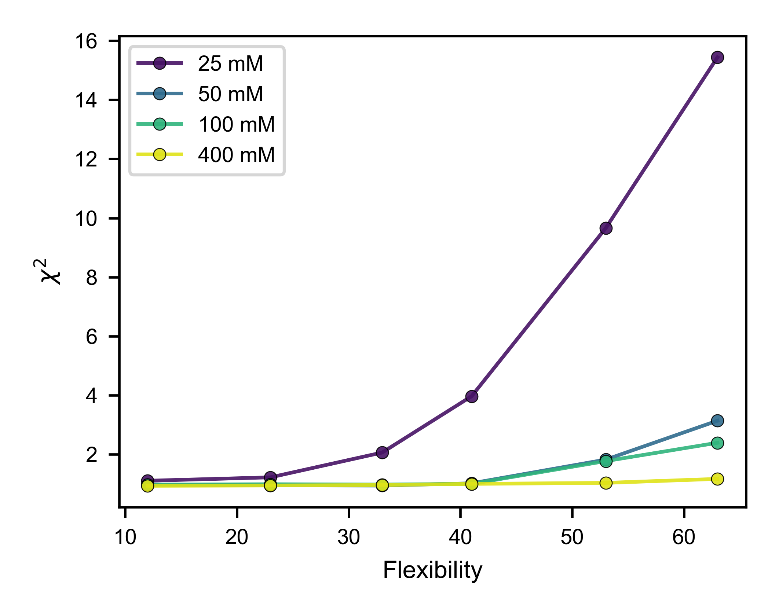

**Figure S3. SAXS ensembles of Phd at varying salt concentrations.** Agreement between the experimental and calculated SAXS curves (χ^2^) for the ensembles of PhD monomers and dimers at varying degrees of dimer unfolding, refined against the experimental SAXS data at different salt concentrations. The ensemble contains a dimer variant with lowest χ^2^ is then shown in Figure 5.
